# Mapping-based all-RNA-information sequencing analysis (ARIseq) pipeline simultaneously revealed taxonomic composition, gene expression, and their correlation in an acidic stream ecosystem

**DOI:** 10.1101/159293

**Authors:** Arisa Tsuboi, Misao Itoga, Yuichi Hongoh, Shigeharu Moriya

**Author notes:** Corresponding author (SM).

## Abstract

We developed a new pipeline for simultaneous analyses of both rRNA profile as a taxonomic marker and mRNA profile as a functional marker, to understand microbial ecosystems in natural environments. Our pipeline, named All-RNA-Information sequencing (ARIseq), comprises a high-throughput sequencing of reverse transcribed total RNA and several widely used computational tools, and generates quantitatively reliable information on both community structures and gene expression patterns, which were verified by quantitative PCR analyses in this study. Particularly, correlation network analysis in the pipeline can reveal microbial taxa and expressed genes that share patterns of dynamics among different time and/or geographical points. The pipeline is primarily mapping-based, using a public database for small subunit rRNA genes and obtained contigs as the reference database for protein-coding genes. We applied this pipeline to biofilm samples, as examples, collected from an acidic spring water stream in the Chyatsubomi-goke Park in Gunma prefecture, Japan. Our analyses revealed the predominance of iron and sulfur-oxidizing bacteria and *Pinnularia* diatoms, and also indicated that the distributions of the iron-sulfur-oxidizing bacterial consortium and the *Pinnularia* diatoms largely overlapped but showed distinct patterns. In addition, our analyses showed that the iron-oxidizing bacterial genus *Acidithiobacillus* and co-occurring *Acidiphilium* shared similar distribution pattern whereas another iron-oxidizing genus *Leptospirillum* exhibited a distinct pattern. Our pipeline enables researchers to more easily capture the outline of microbial ecosystems based on the taxonomic composition, protein-coding gene expression, and their correlations.

## Introduction

Microbial communities play various and crucial roles in their habitat ecosystems. Because most microbial species in many environments are yet uncultivable, culture-independent approaches are generally adopted to reveal both taxonomic compositions and potential functions. Sequencing analysis of small subunit (SSU) rRNA gene amplicons is widely used methodology to obtain information on the taxonomic composition of a microbial community. Metagenome and metatranscriptome analyses are also performed to investigate the community structure based on rRNA and/or other taxonomic marker genes as well as to obtain the functional information on the microbial community. In general, researchers choose a combination of these methodologies to comprehensively understand a microbial ecosystem.

Among these currently available methodologies, RNA-based analyses are essential to the assessment of microbial activities. The large proportion of rRNA, compared to mRNA, has been an obstacle to functional analyses based on mRNA sequences; however, high-throughput sequencing has enabled researchers to analyze sufficient mRNA sequences without depleting rRNA in many cases. Simultaneous acquisition of both mRNA sequences as functional markers and rRNA sequences as taxonomic markers from the same RNA sample is advantageous to depict the precise picture of a microbial ecosystem [1–4]. Furthermore, amplicon-based analyses of rRNA sequences are subjected to PCR amplification bias [5]. Thus, development of a simple and quantitatively reliable pipeline for the simultaneous analysis of both mRNA and rRNA data will greatly help researchers.

Here, we developed a pipeline comprising several steps: a standard quality filtering of sequence reads, mapping reads to a public SSU rRNA sequence database, *de novo* assembly of mRNA reads and annotation of the contigs, mapping reads to the mRNA contigs, and finally statistical analyses of the expression level of respective gene categories and the frequency of microbial taxa. Our pipeline, named All-RNA-Information sequencing (ARIseq), yields quantitatively reliable information for both microbial community structure and gene expression, which was tested by quantitative PCR (qPCR) analyses. The quantitative information is also used for a correlation network analysis, which can reveal microbial taxa and expressed genes that share patterns of dynamics among different time and/or geographical points.

We applied this newly developed tool, ARIseq, to an analysis of biofilm samples collected from an acidic spring water stream at the Chyatsubomi-goke Park in Japan. This acidic stream is a well-known ecosystem for bio-mineralization by iron-oxidizing microbial consortia [6]. ARIseq successfully provided quantitative information on both the microbial community structure and gene expression. The results showed that the similarity and dissimilarity of distribution patterns among dominant organisms and expressed genes.

## Materials and Methods

### Sample collection

Biofilm samples were collected at three different points along an acidic stream at the Chyatsubomi-goke Park in Gunma prefecture, Japan [6]. The stream originated from an acidic spring and then flowed down to the inlet point of the Shirakinu waterfall. The three points were: site 1 upstream point (36°38’58.14"N 138°35’10.66"E), site 2 upstream point (36°38’58.14"N 138°35’11.24"E), and site 3 downstream point (36°38’57.05"N 138°35’27.82"E). A schematic drawing of these sampling sites with water temperature, pH, and ^57^Fe element concentration was shown in Fig 1. The values of pH and water temperature were directly measured by a portable pH meter, HM-30P (TOA DDK, Tokyo, Japan). The total iron concentration in the stream water was determined for ^57^Fe element in the diluted solution by 0.01 mol L^-1^ hydrochloric acid (088-02265, WAKO, Japan) by an inductive coupled plasma mass spectrometry, NexION300 (PerkinElmer Japan, Yokohama, Japan).

**Fig 1.**
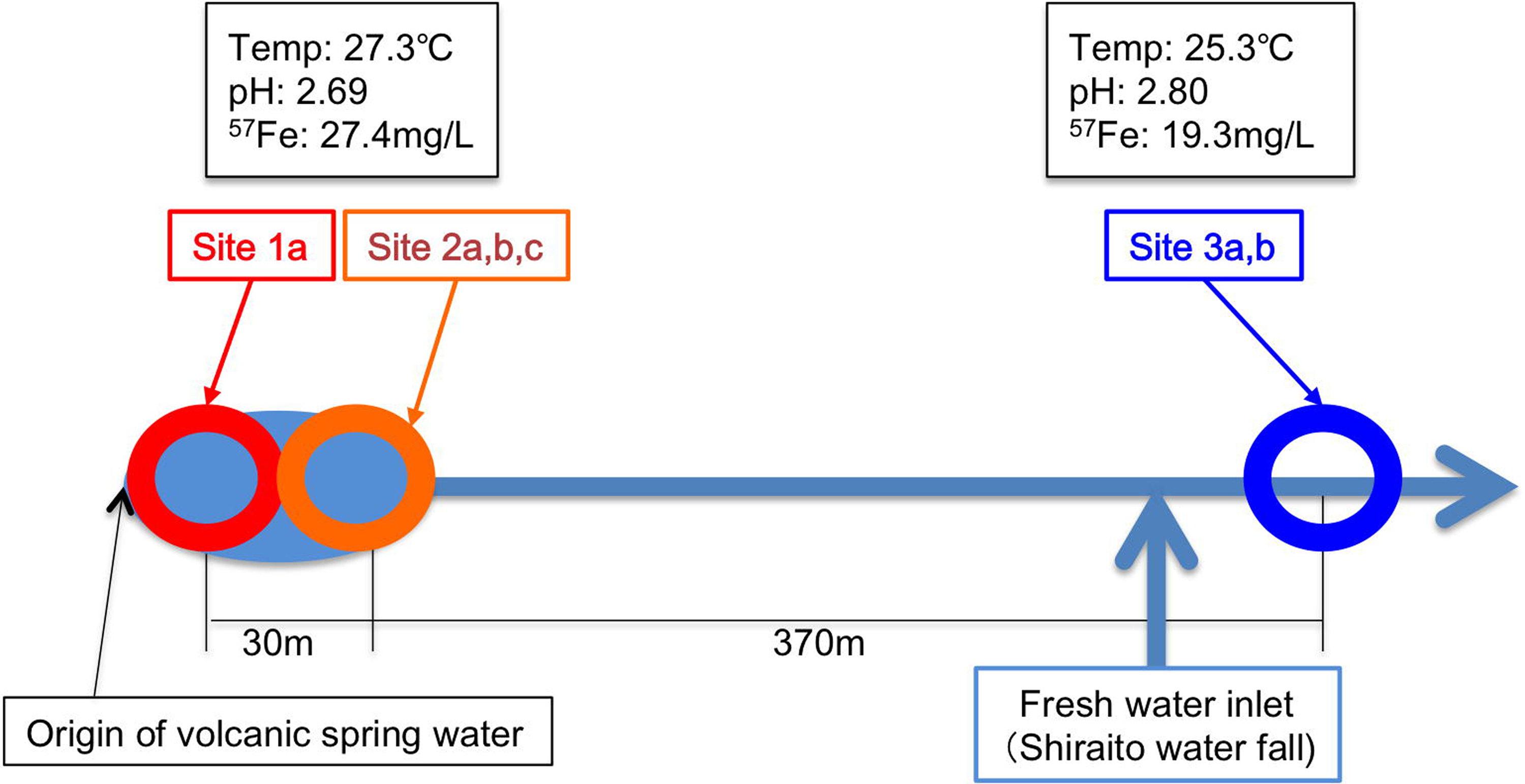
Schematic drawing of sampling site topology and physical and chemical conditions.

Three stones (indicated as a, b, c) per site were sampled. Each stone was rinsed with purified water, and biofilm was brushed off from the stone surface using a sterile toothbrush and purified water. Suspension of the collected biofilm was filtered with a 0.22-μm polyvinylidene difluoride (PVDF) Durapore^®^ membrane (Millipore, Billerica, MA, USA). The filter membrane with trapped materials was preserved in 1 ml of RNA*later*^®^ solution (Ambion, Austin, TX, USA) at -80°C until being processed. In total, nine samples were collected (designated as samples 1a–c, 2a–c, and 3a–c).

### RNA extraction and sequencing

The materials trapped on the filter membrane was resuspended in RNA*later*^®^ by pipetting, and precipitated by centrifugation at 15,000 rpm for 5 min. The pellet was subjected to RNA extraction, using the PowerBiofilm RNA Isolation Kit (MO Bio, Carlsbad, CA, USA), according to the manufacturer’s instructions. RNA was eluted with 100 μl of purified water. RNA concentration was measured using a Qubit system (Invitrogen, Carlsbad, CA, USA) and adjusted to 100 ng/μl with purified water. RNA was not sufficiently recovered from samples 1b, 1c, and 3c, which were removed from further experiments.

Sequencing libraries were prepared using the SMARTer^®^ Stranded RNA-Seq Kit for Illumina (Takara, Kyoto, Japan), following the manufacturer’s instructions. The concentration and length of DNA fragments in the sequence library were measured using the Qubit system and a Bioanalyzer 2100 (Agilent Technologies, Carlsbad, CA, USA). When libraries contained DNA fragments < 80 bp, the libraries were further processed with the Agencourt AMpure XP (Beckman Coulter, Brea, CA, USA) to remove small fragments, according to the manufacturer’s instructions. Sequencing was performed on the Illumina MiSeq platform (Illumina, CA, USA) with the Reagent Kit v3 (600 cycles, paired-end mode). Sequence data have been deposited at DDBJ with the accession number DRA005571.

### Processing sequence data

The scheme of sequence processing is outlined in Fig. 2. Sequence reads were trimmed and quality-filtered using program Trimmomatic [7] in paired-end mode with a seed mismatch value of 5, a palindrome clip threshold of 30, a simple clip threshold of 7, a minimum read length of 100 bp and a headcrop of 6 bp. Trimmed reads were used for mapping to the SSU rRNA sequence database SILVA release 108 for QIIME [8,9]. Mapping was performed using Bowtie2 [10] with a local alignment mode and single- or paired-end modes. The data generated by Bowtie2 were converted to the sorted binary sequence alignment/map (BAM) format, using samtools [11], and the number of mapped reads were counted using the eXpress program package [12]. The read count data were integrated into taxon information and utilized for secondary analyses (see below).

**Fig 2.**
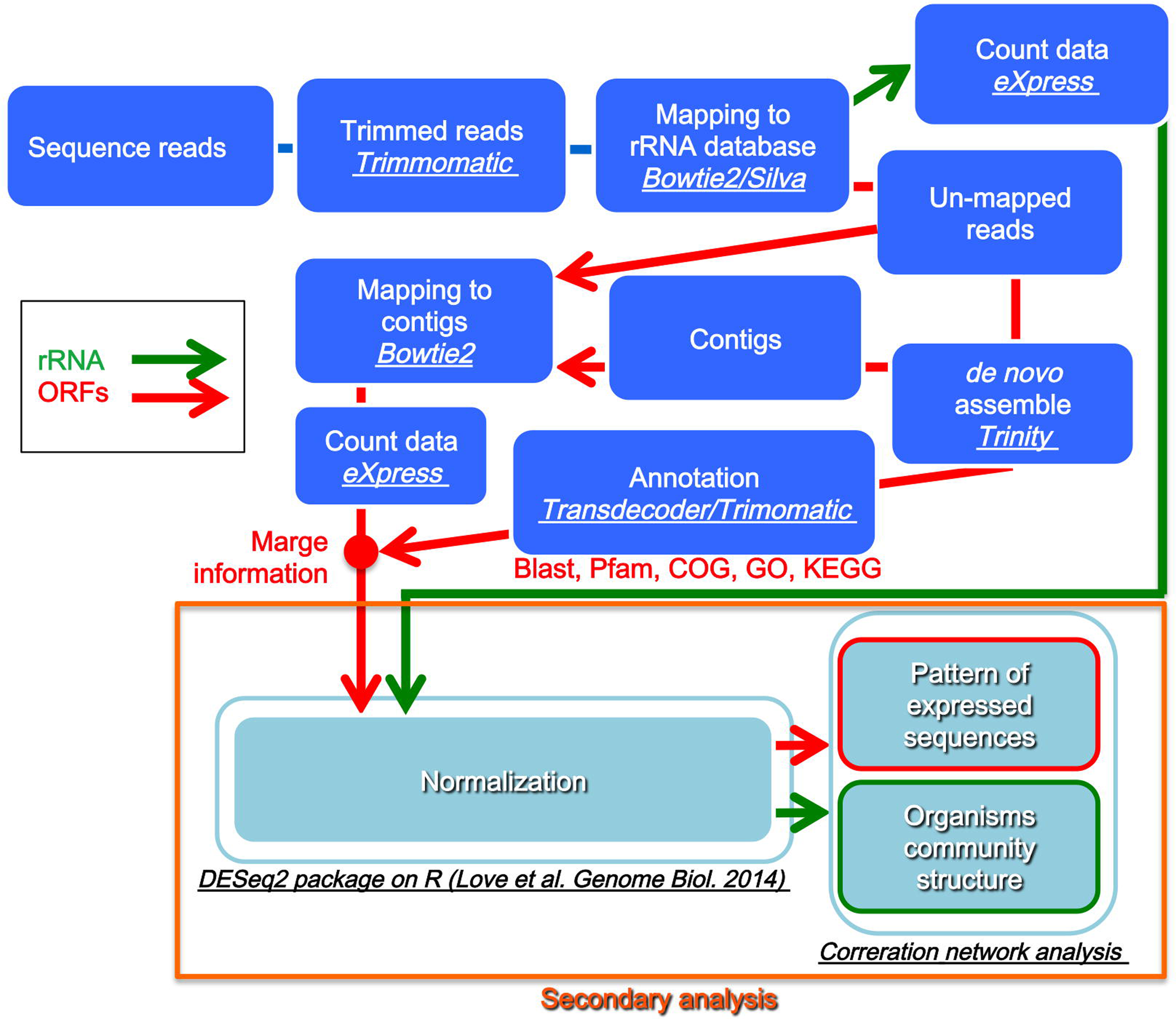
Schematic flow chart of the ARIseq pipeline. Green arrows indicate the flow of rRNA gene data, and red arrows indicate the flow of the other RNA.

Reads unmapped to the SILVA SSU rRNA database by paired-end mode were assembled using the Trinity program package [13] with the Jaccard clip option, to construct contigs. Open reading frames (ORFs) and the encoded protein sequences were predicted using the TransDecoder program package (https://transdecoder.github.io/). The ORF data were used as the reference database for mapping reads. Functional annotation of the identified ORFs was conducted with the Trinotate program package (https://trinotate.github.io/) that uses a combination of a BlastP search against the “UniProt/Swiss-Prot database for Trinotate”, an hmmer search in the Pfam database, and RNAMMER analysis. BlastP searches against the non-redundant (nr) and standard Swiss-Prot/UniProtKB databases were conducted, and the results were added to the annotation. Furthermore, a Ghost KOALA search provided by the Kyoto Encyclopedia of Genes and Genomes (KEGG) [14] was performed, and a K number was assigned to each of the ORFs. These functional annotations were combined with the read count data, and rank abundance curves were constructed.

### Secondary analysis

The read count data for both SSU rRNA and ORFs were subjected to secondary analyses. First, we performed normalization of the read count data, following a negative binomial distribution with generalized linear model, using the DESeq2 package [15]. For this analysis, the reads were divided into two groups: the “upstream group” (samples 1a and 2a–c) and the “downstream group” (samples 3a, b). SSU rRNA reads or ORFs with less than 10 mapped reads in total from all samples were excluded. The calculation with the DESeq2 package was performed using the TCC package in *R* [15].

Network analysis was performed based on Spearman’s rank correlation coefficient matrix, calculated using *R*, for the normalized read count datasets. The results were visualized using the Gephi program package [16] with a transformed matrix of connection between source and target with collected high positive correlation coefficient (*r* > 0.7) from correlation coefficient matrix [17]. The raw read count datasets were used also for calculating rarefaction curves using Past3.14 [18].

### Quantitative PCR experiments

We performed qPCR to quantify the expression level of genes and to compare the results with those obtained from our pipeline. We selected two SSU rRNA sequences and two contigs each containing an ORF (ORF contig) as examples that showed differential expression patterns. We chose taxon 17056 (18S rRNA of *Pinnularia* cf. *gibba)* and the ORF contig TRINITY_DN8038_c26_g13_i2|m.9499 (peptide 9499), as they were highly expressed in the “upstream group”, and taxon 50111 (18S rRNA of *Chironomus tentans*) and the ORF contig TRINITY_DN6923_c3_g1_i3|m.6571 (peptide 6571), as they were highly expressed in the “downstream group”. Peptide 9499 was annotated as an “uncharacterized protein” by both the Trinotate pipeline and a BlastP search against the Swiss-Prot/UniProtKB database. Peptide 6571 was predicted to be 3-dehydroquinate dehydratase/shikimate dehydrogenase (K13832) by Ghost KOALA or putative alpha-L1 nicotinic acetyl choline receptor by a BlastP search against the Swiss-Prot/UniProtKB database. Primer sequences for qPCR are shown in Table. 1.

Sequencing libraries were used as templates for qPCR, although one of the downstream samples (sample 3b) was excluded because the library DNA ran out by sequencing. Four samples (1a and 2a–c) from the upstream group and one (3a) from the downstream group were subjected to qPCR using the KAPA^™^ SYBR^®^ Fast qPCR Kit (KAPA Biosystems, MA USA), according to the manufacturer’s instructions. The qPCR was performed in a Thermal Cycler Dice^®^ Real Time System TP850 (Takara) in 25-μl reaction volume, and the PCR program was as follows: initial denaturation at 95°C for 30 sec and 40 cycles of 95°C for 30 sec and 60°C for 1 min. The dilution rate of the samples was 1/200. The calibration curve was constructed using sample 2c with dilution rates 1/100, 1/200, 1/400, 1/800, 1/1600, and 1/3200. The experiments were conducted in triplicate. The amount of template DNA was adjusted, using the KAPA^™^ Library Quantification Kit for Illumina (KAPA Biosystems, MA USA).

## Results and Discussion

### Mapping-based analysis of total RNA sequences

Total RNA sequencing of the six biofilm samples (1a, 2a–c, and 3b,c) generated 770,334 to 1,320,217 read pairs, and >80% reads passed the quality check (Table 2). These reads were analyzed using our mapping-based ARIseq pipeline (Fig 2). Of the reads, 45.7% were mapped to the SILVA SSU rRNA sequence database, using the single-end mode option of Bowtie2 (Table 2). When using the paired-end mode, mapped reads were only 5.1%; thus, we used the results obtained using the single-end mode for quantification data. We here only considered quantification of SSU rRNA genes sharing a high sequence identity with those in the reference database, Therefore, the unmapped reads should contain other SSU rRNA, large subunit rRNA, and non-coding RNA, which can be identified and removed in the following annotation step.

Unmapped reads against the SILVA database by paired-end mode mapping were used for a *de novo* assembly process using Trinity, and then ORFs on the contigs were predicted by using TransDecoder. These ORFs were used as the reference sequence database for mapping analysis to obtain gene expression profiles. In the single-end mode of Bowtie2, 17.2% of the reads were mapped onto this self-made database, whereas only 2.7% of the reads were mapped in the paired-end mode, (Table 2). Here, we again employed the results from the single-end mode for subsequent analyses.

Read count data for both SSU rRNA and ORFs were obtained using the eXpress program package. Fig 3 shows rarefaction curves, which indicated that the sequence efforts (*i.e.,* number of reads) for both rRNA and ORFs were sufficient or nearly sufficient to obtain most variations in the RNA samples except for the SSU rRNA of the downstream samples 3a and 3b. All read count datasets were subjected to the normalization processes with DESeq2 and annotation using Trinotate, BlastP, and Ghost KOALA (see Materials and Methods for details). The annotation and normalization results were listed in S1 and S2 Tables.

**Fig 3.**
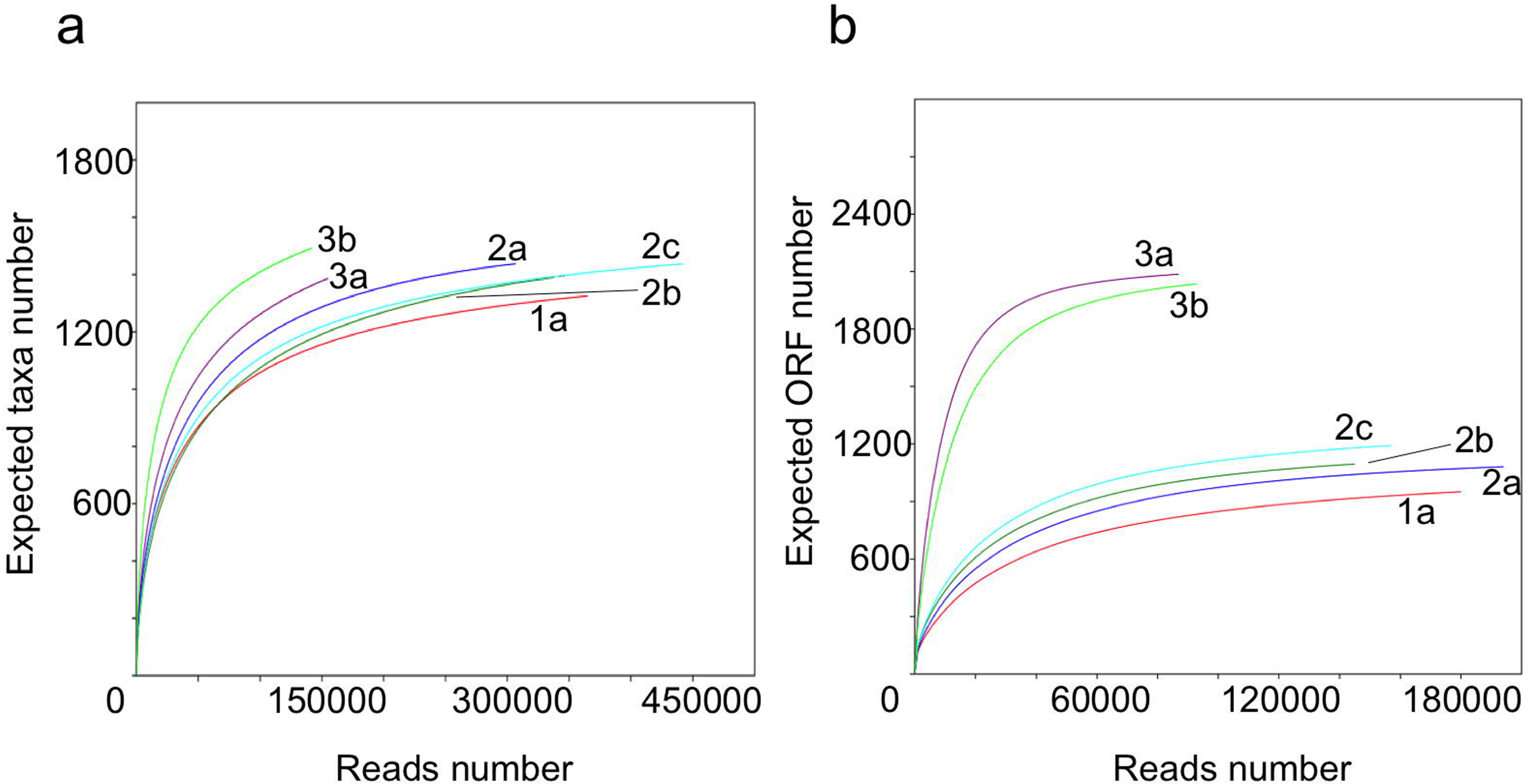
Rarefaction curves. (a) Rarefaction curves of taxa based on SSU rRNA sequences. Taxa was defined as reads mapped OTUs. (b) Rarefaction curves of protein-coding genes (ORF) based on cDNA sequences from mRNA. Sample names are shown aside the curves.

In the normalized read count data, gene expressions of several rRNA or ORFs greatly differed between the upstream and the downstream sample groups. For example, mapping data suggested that, a diatom, *Pinnularia* cf. *gibba* (taxon 17056), was abundant in the upstream samples but less represented in the downstream sample 3a based on the rRNA read count data (Fig 4a). In contrast, aquatic larvae of the non-biting midge *Chironomus tentans* (taxon 5111) were found abundant only in the downstream sample 3a (Fig 4a). Although insects are not microbes, we here included them because they probably have a great impact on the biofilm ecosystem. These expression patterns were well congruent with the results obtained by qPCR analyses (Fig 4b). ORF read count data showed that the ORF for peptide 9499 was most highly expressed in the upstream samples, whereas ORFs for peptide 6571 was most highly expressed in the downstream samples (Fig 5a). These results were also congruent with those from qPCR analyses (Fig 5b). Thus, our ARIseq analysis pipeline generated quantitatively reliable results for both rRNA and expressed ORFs.

**Fig 4.**
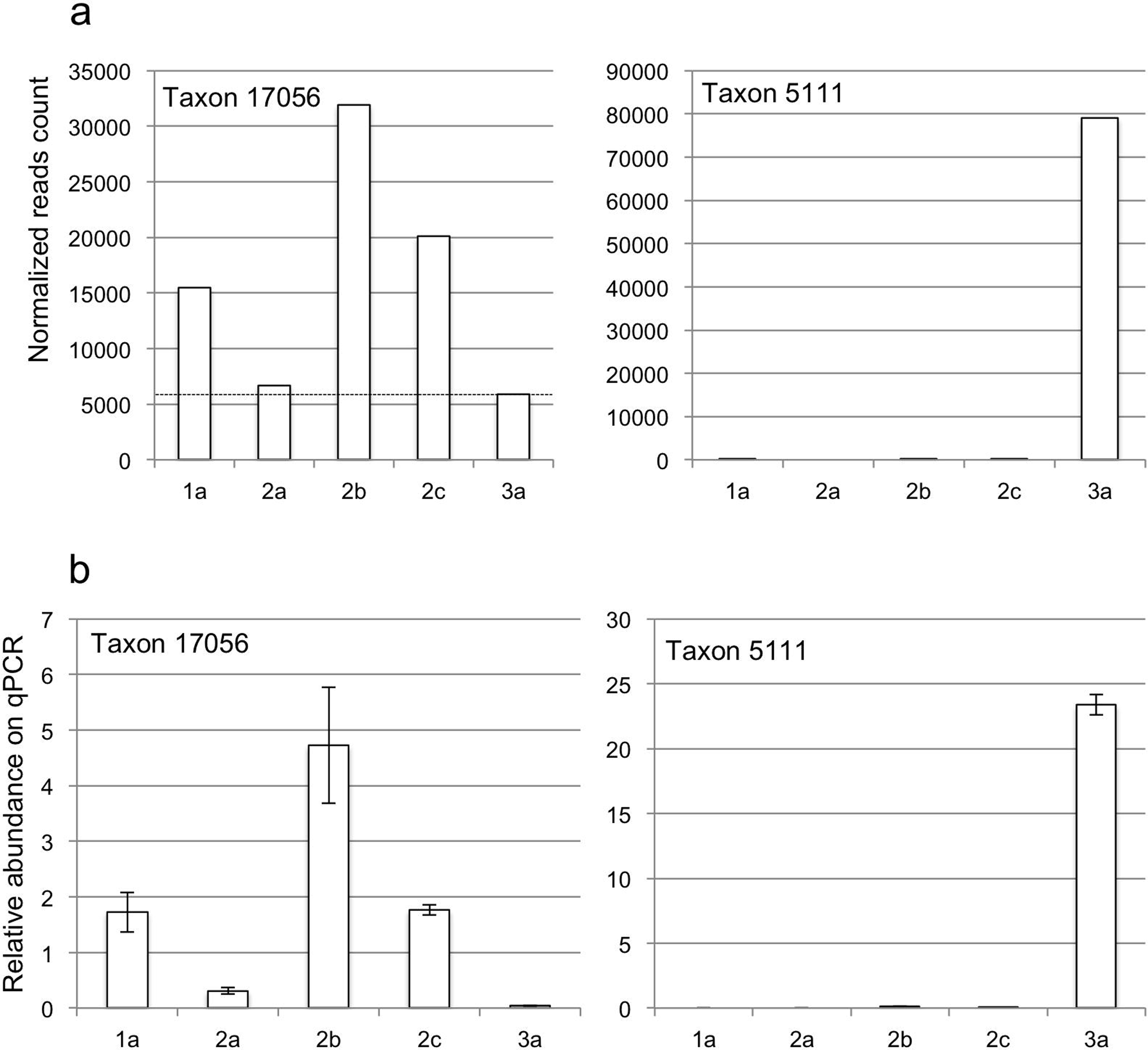
Quantitative analysis of rRNA data. (a) Normalized read counts for an upstream-specific taxon, 17056, and a downstream-specific taxon, 5111. (b) Relative abundance of these taxa evaluated by qPCR.

**Fig 5.**
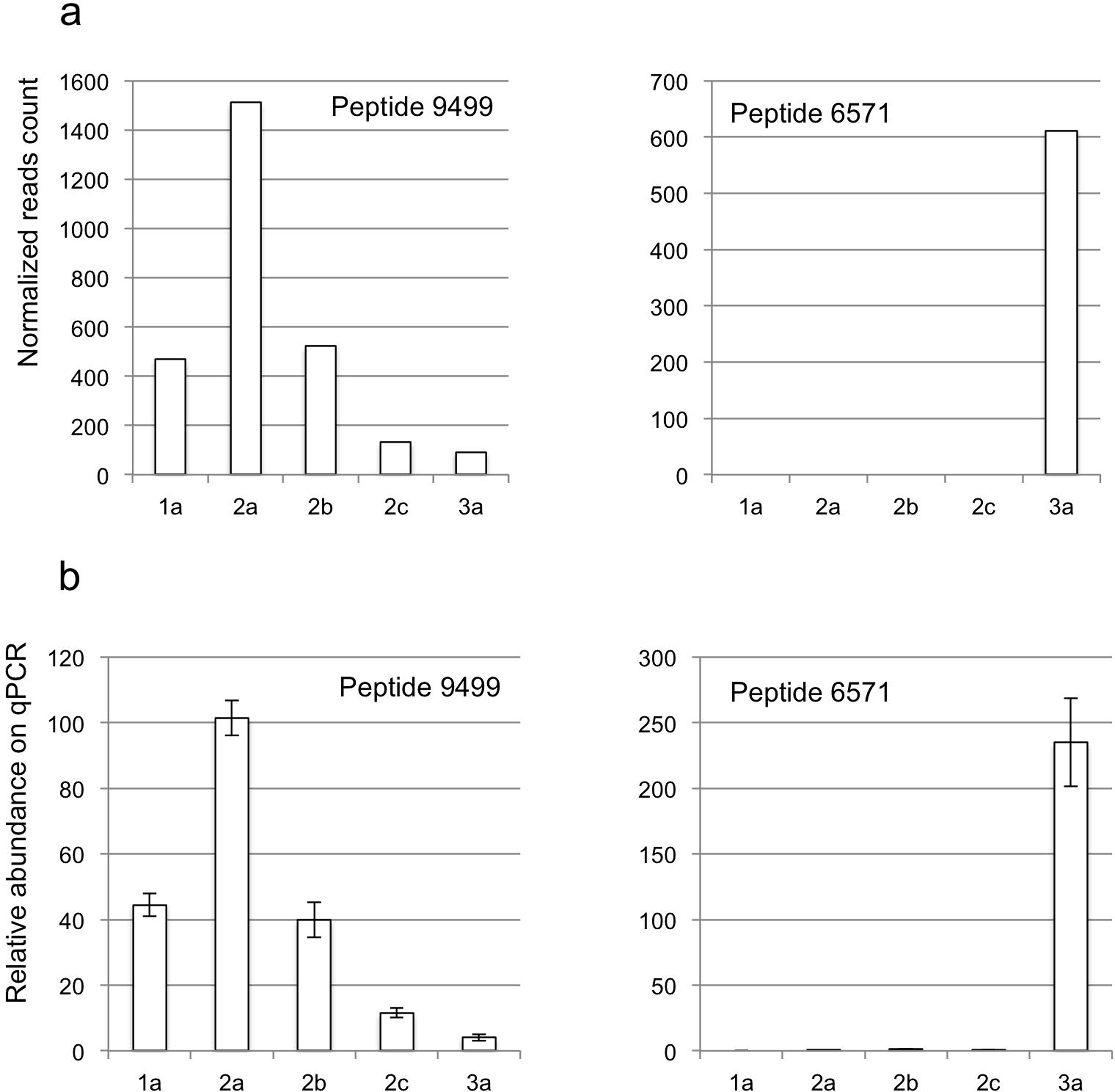
Quantitative analysis of ORF data. (a) Normalized read counts of an upstream-specific ORF for peptide 9499 and a downstream-specific ORF, 6571. (b) Relative abundance of these ORFs evaluated by qPCR.

### Biofilm community structure in the acidic stream

Fig 6 shows SSU rRNA-based taxonomic compositions at the domain level. Eukaryotes or bacteria predominated in all samples; only a few archaeal sequences were detected. The distribution pattern of each taxon was summarized by a correlation network analysis (Fig 7a). Six clusters were recognized, and each distribution pattern is shown in Fig 7b. Here, a “cluster” is defined as a group of taxa that exhibit similar distribution pattern among the samples. Members of cluster 1 mainly inhabited the downstream sites (samples 3a,b), whereas members of clusters 3, 4, and 5 were found mostly at upstream sites (samples 1a and 2a–c). Because the reads assigned to clusters 2, 5, and 6 were relatively few, we focused only on clusters 1, 3, and 4 for comparisons.

**Fig 6.**
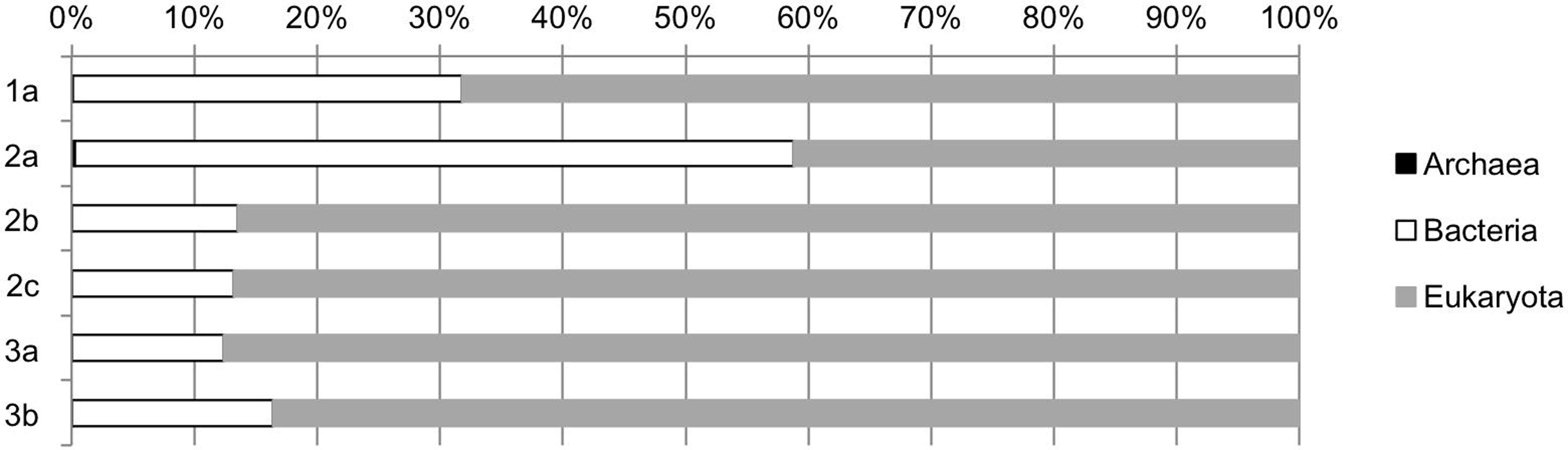
Taxonomic comopistions at domain level. Normalized read counts are used for the calculation.

**Fig 7.**
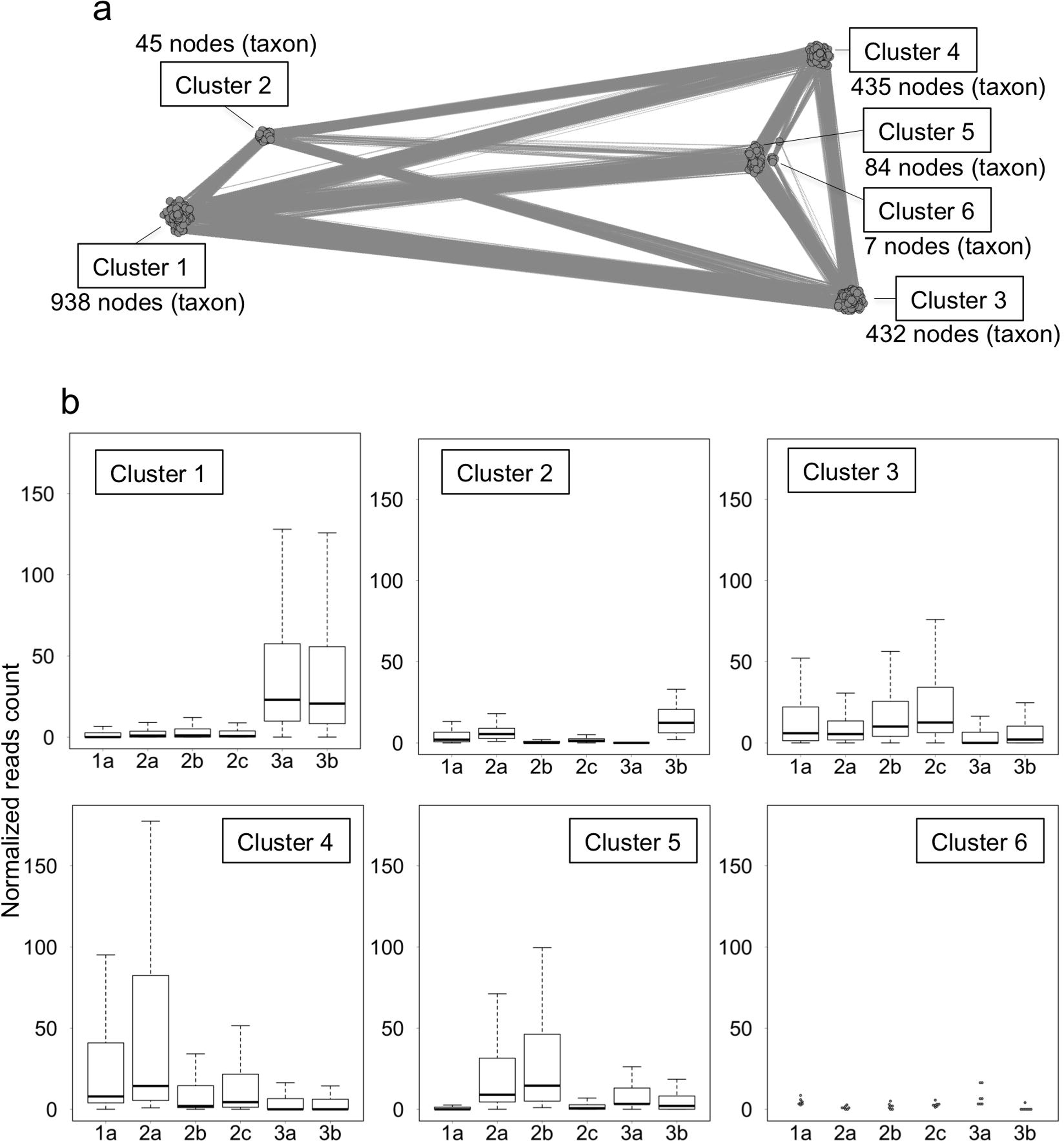
Correlation network analysis of SSU rRNA data. (a) A network drawn for relationships with positive correlation indexes > 0.7. (b) Normalized read counts of SSU rRNA for taxa assigned to each cluster.

Fig 8 shows the taxonomic composition of each cluster at the genus level. *Chironomus* midge larvae, *Pinnularia* diatoms, and *Acidithiobacillus* bacteria predominated in clusters 1, 3, and 4, respectively. This at once indicated that *Chironomus* predominated in the downstream regions, and that *Pinnularia* and *Acidithiobacillus* did in the upstream regions. *Acidithiobacillus* is a well-known acidophilic bacterial genus that oxidizes sulfur and iron, and is frequently found in acid mine drainage [19–22]. In this acidic stream, the presence of bacteria attached to iron precipitate with phosphorous/sulfur crystals was previously reported [6]. It is highly possible that *Acidithiobacillus* members with co-occurring, possibly symbiotic *Acidiphilium* [23], the second-dominant genus in cluster 4, mainly cause iron/sulfur oxidation.

**Fig 8.**
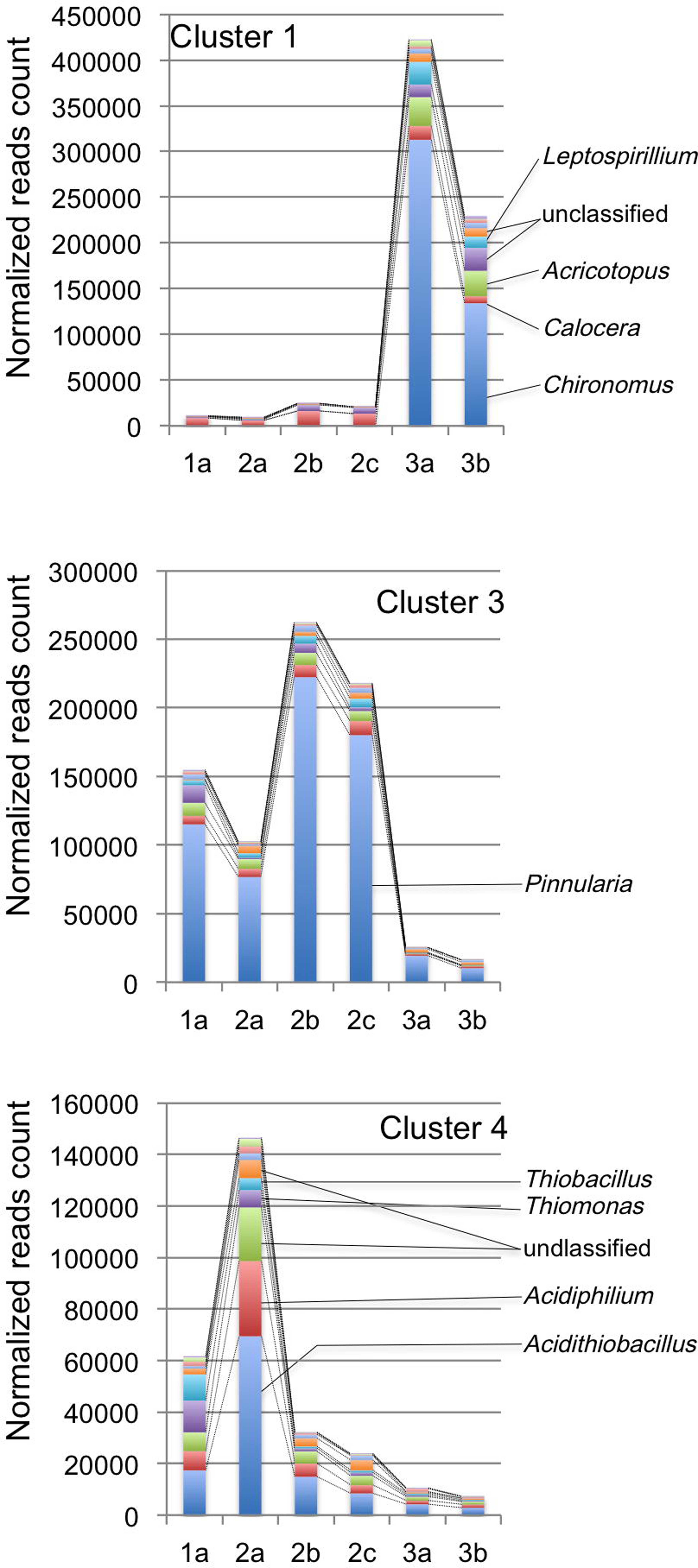
Taxonomic compositions of each cluster generated by network analysis of SSU rRNA data. Normalized read counts are shown at the genus level. The taxonomy was based on the SILVA database release 108. Original data are shown in S3 Table.

*Pinnularia* diatoms are also frequently found in acid mine drainage [19,24] and comprise a biofilm community [25]. The presence of *Pinnularia-like* diatoms in this acidic stream was previously reported: the cells were found around or attached to precipitated Fe-P materials [6]. As shown in Fig 8, overlapping but distinct distribution pattern is observed between *Pinnularia* and *Acidithiobacillus* in the upstream region. This pattern suggested that the primary production at the upstream site 2a is largely attributable to the iron-sulfur oxidation by *Acidithiobacillus,* while photosynthesis by *Pinnularia* contributes to the primary production broadly in the upstream region

In the downstream samples, the larvae of the non-biting midge genera *Chironomus* and *Acricotopus* were predominant (Fig 8). These chironomid larvae are detritivores and frequently found in biofilms in freshwater [26]. The third-dominant genus in the downstream samples in cluster 1 was *Leptospirillum*. *Leptospirillum* members are also iron-oxidizers in general and produce iron/sulfur granules inside their extracellular polymeric substances that compose the biofilm [19,27,28]. This different distribution pattern between *Acidithiobacillus* and *Leptospirillum* as revealed in our correlation analysis implies their niche differentiation.

### Gene expression profiles and correlation with taxonomic compositions

ORFs showing a similar expression pattern among the samples were also clustered by a correlation network analysis (Fig 9a). Two large clusters were generated: cluster 1 comprised ORFs that were highly expressed in the downstream samples; cluster 2 comprised those highly expressed in the upstream samples (Fig 9b). In cluster 2, genes related to photosynthesis, such as electron transportation and photosystem components were dominant (Table 3). This coincided with the predominance of diatoms including *Pinnularia* in the upstream region (Fig 8).

**Fig 9.**
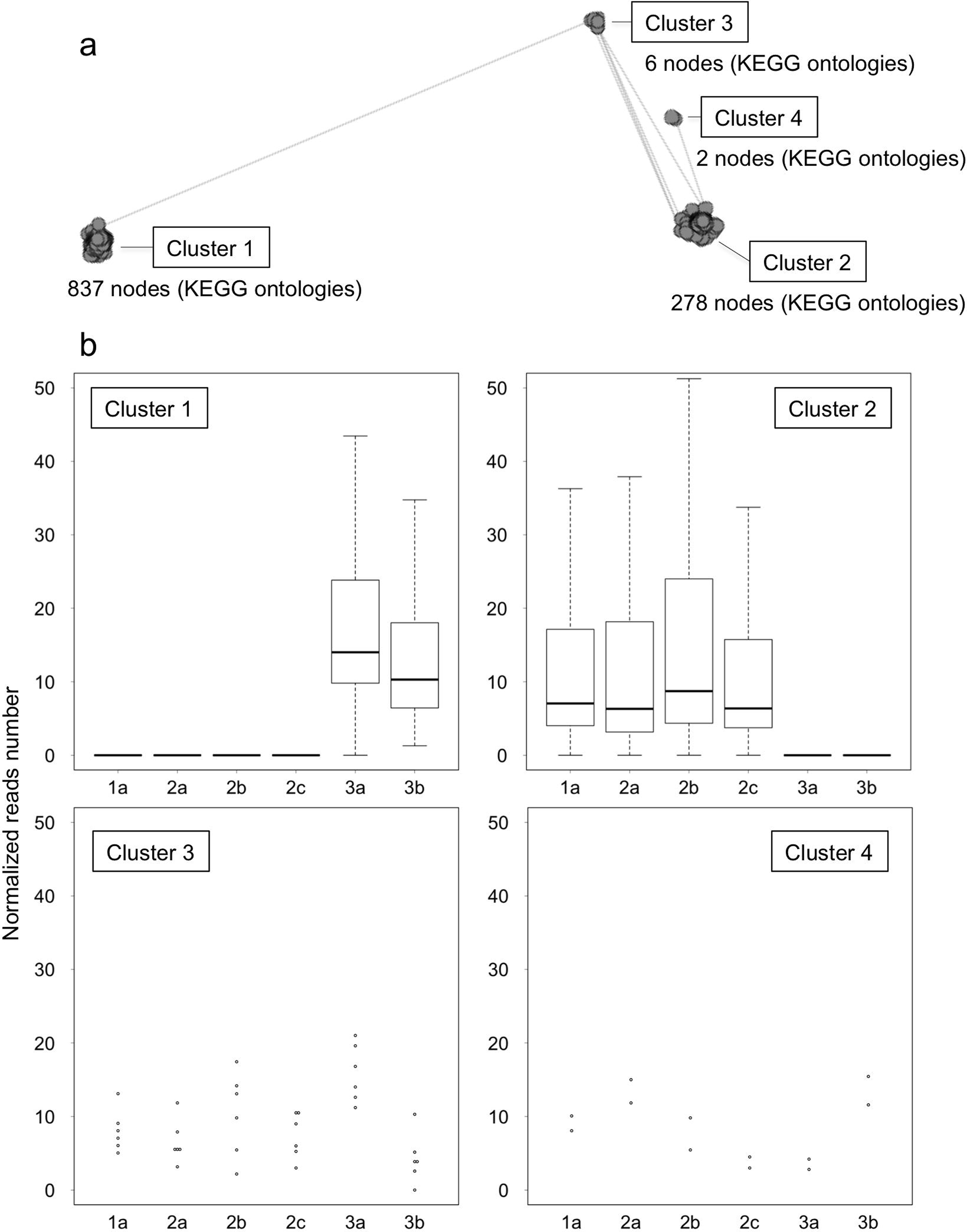
Correlation network analysis of expressed ORF data. ORFs were categorized and bundled according to the KEGG orthology. Normalized read counts were used for the analysis. (a) A network drawn for relationships with positive correlation indexes >0.7. (b) Normalized read counts of expressed ORFs assigned to each cluster.

Furthermore, we constructed a correlation network based on expression patterns of both rRNA and ORF datasets (Fig 10). Here, “cluster” comprised both SSU rRNA and ORFs that exhibited similar distribution patterns among the sampling sites. The clustering pattern resembled that based on SSU rRNA (Fig 8 and 11); the ORF expression patterns and the SSU rRNA distribution patterns were well congruent. Dominant gene categories in each cluster were mostly house-keeping genes (Tables 4 and 5) or photosynthesis-related genes (Table 6). Rank abundance curves of taxa that expressed genes related to photosynthesis and assigned to cluster 6 are shown in Fig 12, which suggested that photosynthesis-related genes were mainly originated from diatoms. In Fig 12, closest species that were identified by BlastP searches against the SwissProt/UniProtKB database are shown. *Phaeodactylum tricornutum,* listed as the predominant species in Fig 12, is a marine diatom and one of the few diatoms with genome sequence being analyzed [29]; thus, this most likely represented the predominant diatom *Pinnularia* in the upstream region of this acidic stream.

**Fig 10.**
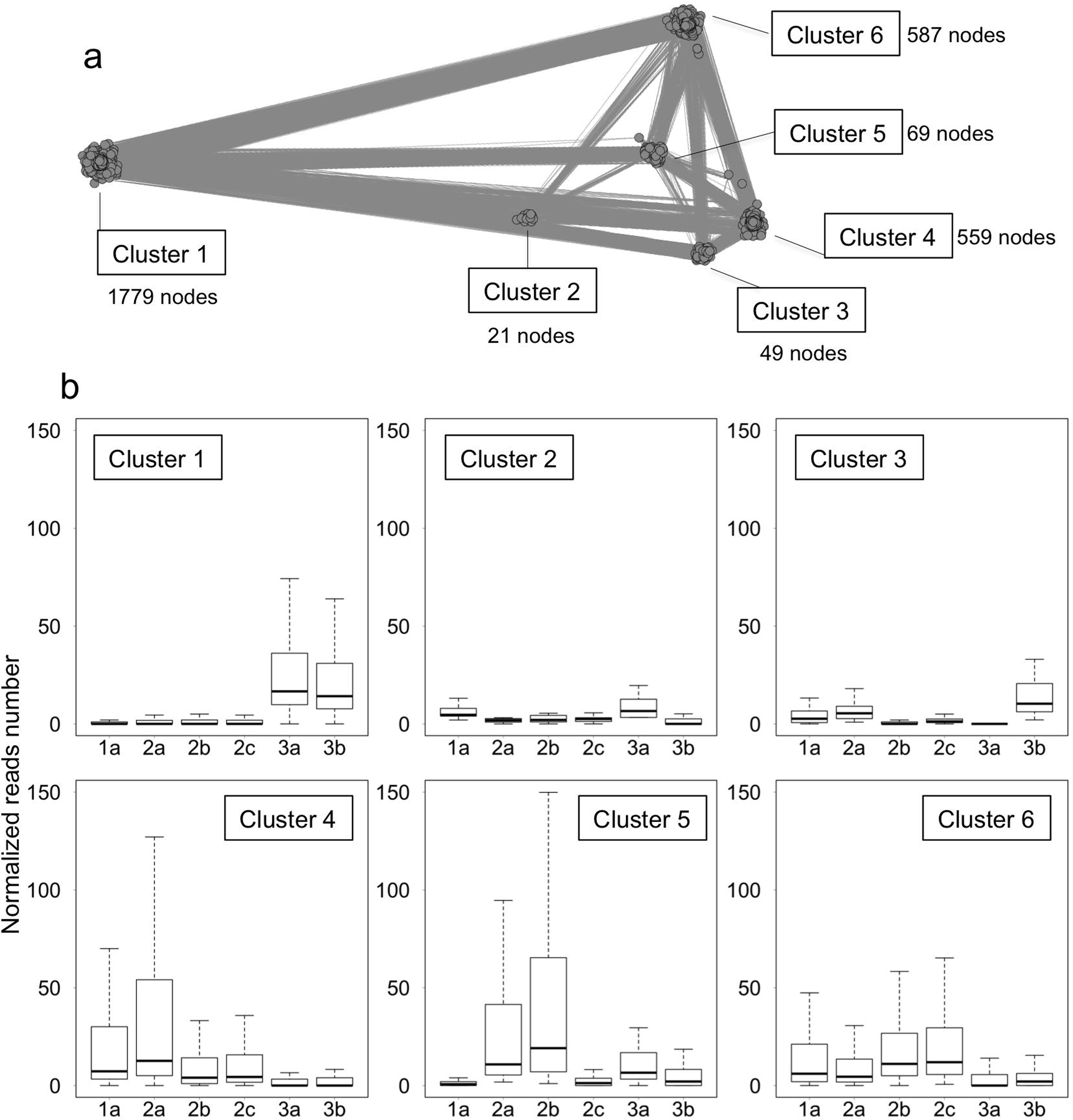
Correlation network analysis of combined data of SSU rRNA and expressed ORFs. (a) A network was drawn for relationships with positive correlation indexes >0.7. (b) Normalized read counts of SSU rRNA-based taxa and expressed ORFs assigned to each cluster.

**Fig 11.**
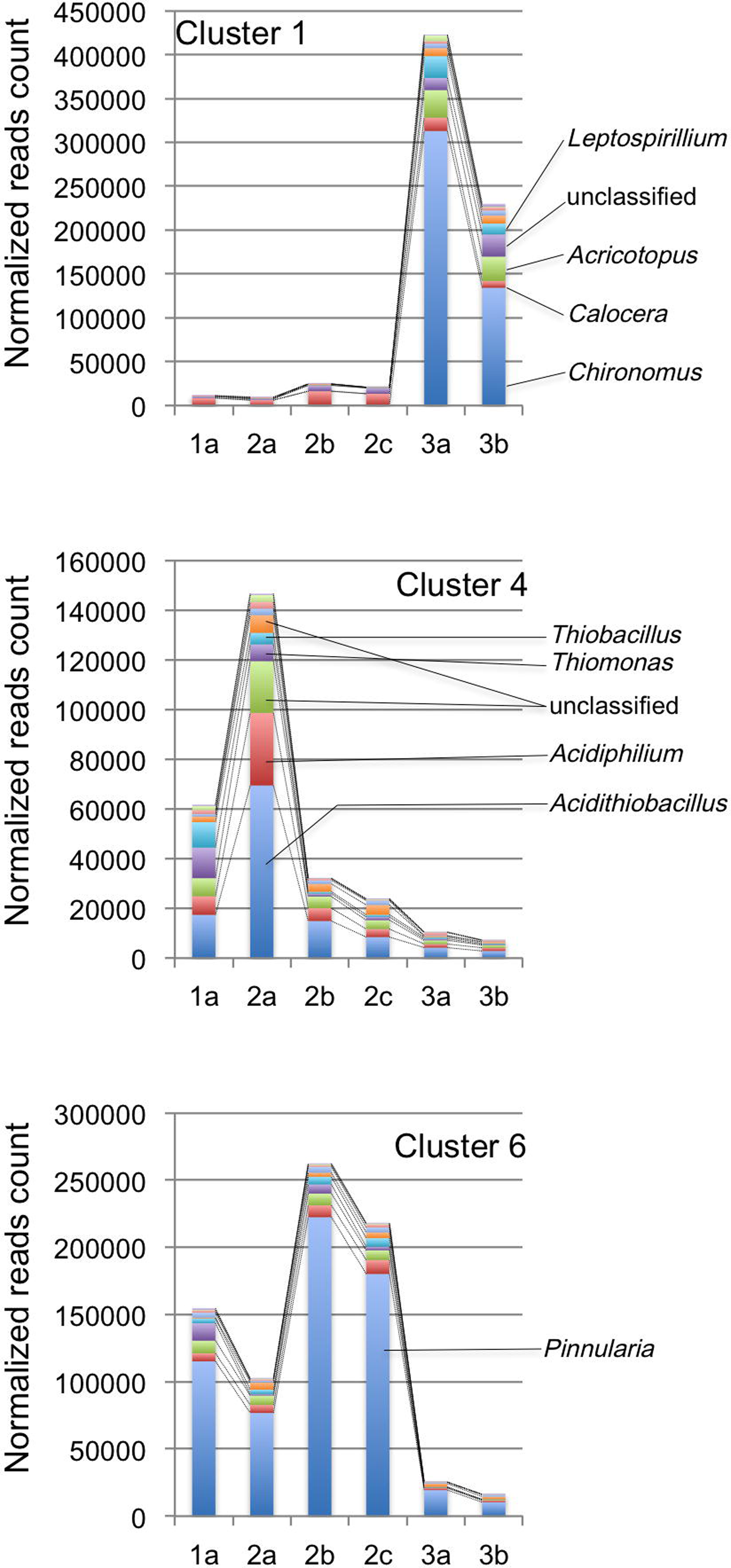
Taxonomic compositions of each cluster generated by network analysis of combined data of SSU rRNA and expressed ORFs. The taxonomic compositions were based only on the SSU rRNA data. Original data are shown in S4 Table. See also the legend to Fig 8.

**Fig 12.**
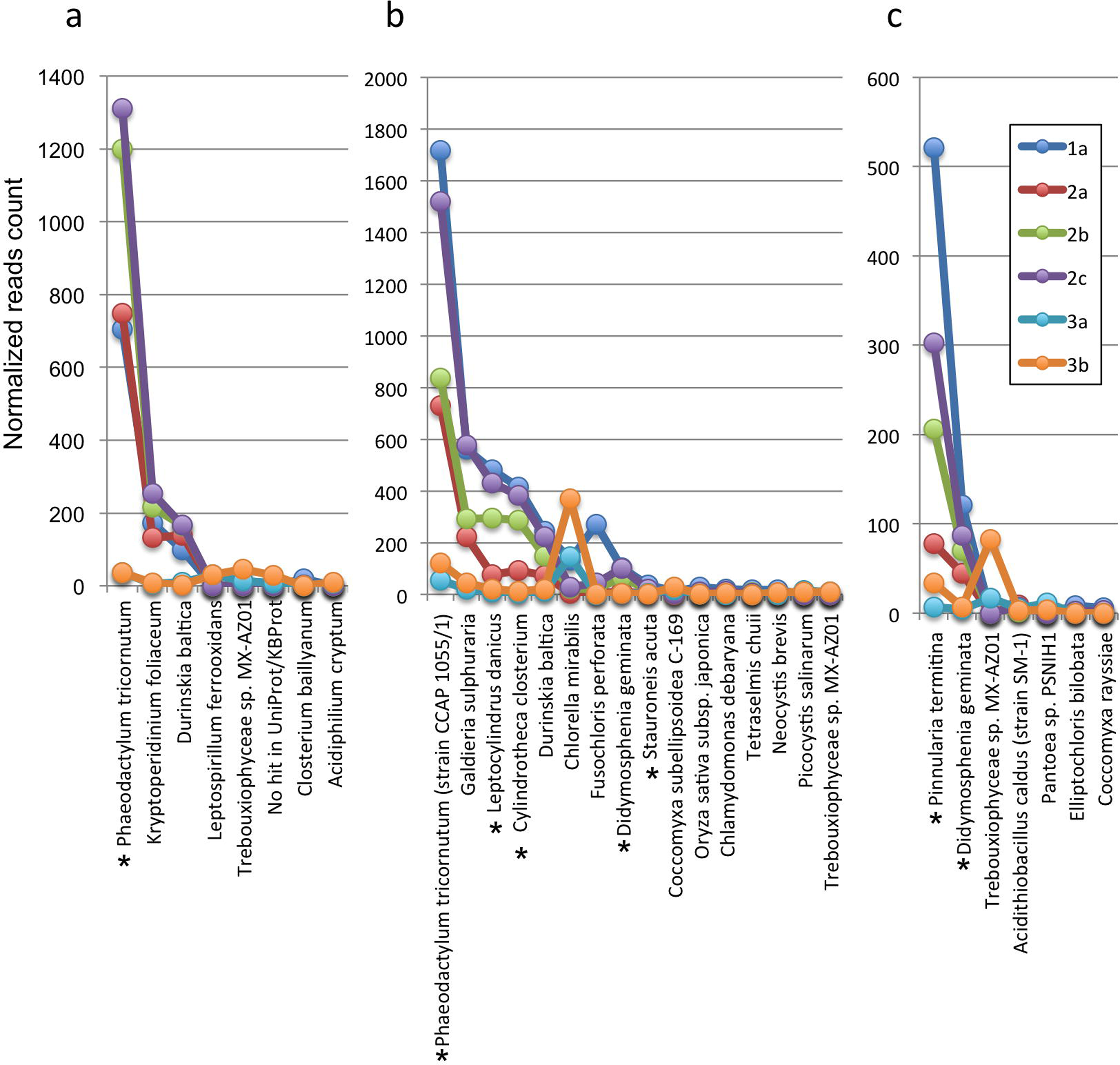
Rank abundance curves of taxa expressing genes involved in photosynthesis.

The species names are the closest taxa found by BlastP searches for photosynthesis-related genes against the SwissProt/UniprotKB database. Genes assigned to cluster 6 in Fig 10 were used for the analysis. (a) Rank abundance curves of organisms that expressed genes related to electron transportation (K00330, K00339, K00343, K00412, K02256, K02261, K02262, K03881, K03883, K03934, and K03935). (b) Rank abundance curves of organisms that expressed genes related to photosystem (K02689, K02690, K02703, K02704, L02705, K02706, and K08910). (c) Rank abundance curves of organisms that expressed genes for the RuBisCO components (K01601 and K01602). Asterisks indicate diatoms.

## Conclusions

Our All-RNA-Information Sequencing analysis (ARIseq) successfully revealed both microbial community structures and expressed gene categories in quantitatively reliable forms. Particularly, our ARIseq pipeline was able to show the correlation among the community members based on rRNA frequency and also the correlation among the gene categories in the biofilm samples collected from the acidic spring water stream. Our pipeline is suitable for researches to capture comprehensive information from one RNA sequencing analysis and facilitates understanding of ecological functions of organismal communities in natural environments.

## Acknowledgements

Sampling was carried out in an environmental research permitted by the board of education, Gunma Prefecture, Japan.

## Contributions

SM, AT, and YH designed the research. AT and MI performed the sampling. AT and SM performed the experiments and analyses. AT, SM, and YH wrote the paper.

## Corresponding author

Shigeharu Moriya

## Supporting information

S1_Table. Annotation of SSU rRNA and data matrix.

S2_Table. Annotation of expressed ORFs and data matrix.

S3_Table. Taxonomic composition of each cluster based on SSU rRNA.

S4_Table. Taxonomic composition of each cluster based on SSU rRNA and expressed ORFs.

